# Deep-mining of vertebrate genomes reveals an unexpected diversity of endogenous viral elements

**DOI:** 10.1101/2023.10.26.564176

**Authors:** Jose Gabriel Nino Barreat, Aris Katzourakis

## Abstract

Endogenous viral elements (EVEs) are key to our understanding of the diversity, host range and evolutionary history of viruses. Given the increasing amounts of virus and host sequence data, a systematic search for EVEs is becoming computationally challenging. We used ElasticBLAST on the Google Cloud Platform to perform a comprehensive search for EVEs (kingdoms *Shotokuvirae* and *Orthornavirae*) across vertebrates. We provide evidence for the first EVEs belonging to the families *Chuviridae*, *Paramyxoviridae*, *Nairoviridae* and *Benyviridae* in vertebrate genomes. We also find an EVE from the *Hepacivirus* genus of flaviviruses with orthology across murine rodents. Phylogenetic analysis of hits closely related to reptarenavirus and filovirus ectodomains suggest three independent captures from a retroviral source. Our findings increase the family-level diversity of non-retroviral EVEs in vertebrates by 44%. In particular, our results shed light on key aspects of the natural history and evolution of viruses in the phyla *Negarnaviricota* and *Kitrinoviricota*.

---

Viruses of all genome types can potentially integrate into host genomes and give rise to endogenous viral elements (EVEs) (1). An EVE forms when viral genetic information enters the host germline and is transmitted vertically to offspring. A novel EVE exists initially as an insertion polymorphism, but can eventually reach fixation subject to the forces of natural selection and genetic drift (1). These fixed EVEs have the highest chance of surviving long periods of time in host genomes, and therefore provide valuable information on virus-host associations over geological timescales. In particular, discovery of endogenous viruses can expand both taxonomic and biogeographical host range, as well as establish direct timelines of association between virus and host (2,3). As such, EVEs constitute a genomic fossil record preserving information on ancient viruses and their interactions.

Although the majority of EVEs in vertebrate genomes are of retroviral origin, non-retroviral EVEs have also been described. Currently, the non-retroviral EVEs found in vertebrates can be assigned to 5 viral kingdoms: *Pararnavirae* (family *Hepadnaviridae*) (4), *Heunggongvirae* (family *Herpesviridae* and *Teratorns*) (5,6), *Bamfordvirae* (*Mavericks*/*Polintons*) (7), *Shotokuvirae* (families *Parvoviridae* and *Circoviridae*) (8,9) and *Orthornavirae* (families *Bornaviridae*, *Filoviridae* and *Flaviviridae*) (10–12). Apart from *Teratorns* and *Mavericks*, other non-retroviral elements found in vertebrate genomes lack self-encoded integrases (5–7). In humans, the herpesvirus HHV6 can integrate a full copy of its genomes into telomeric regions by homologous recombination (13), and these are known to be transmitted vertically (14). EVEs from other viral families tend to be found as fragmentary elements rather than full genomic copies, although full-length EVEs have been reported for hepadnaviruses, circoviruses and parvoviruses (4,9,15,16).

In vertebrates, non-retroviral EVEs from the kingdoms *Shotokuvirae* and *Orthornavirae* are among the most abundant and diverse EVEs. The kingdom *Shotokuvirae* comprises 16 families of ssDNA and dsDNA viruses that descend from an ancestral HUH (histidine-hydrophobic-histidine endonuclease) encoding virus (17,18). The kingdom *Orthornavirae* comprises 112 families of RNA viruses which encode the RNA-dependent RNA polymerase (RdRp) (18). Both shotokuviruses and orthornaviruses include members which are pathogenic to vertebrates. For example, in parrots (Psittacidae), the circovirus Beak and feather disease virus can cause immunosuppression and loss of feathers, with potentially fatal outcomes (19). Canine parvovirus is highly contagious and can cause serious illness in domestic and wild canids (20). Multiple families in the kingdom *Orthornavirae* are known to be highly pathogenic to humans and other vertebrates. Members of the families *Filoviridae*, *Arenaviridae*, and *Nairoviridae* can cause haemorrhagic fevers with high case fatality rates (up to 30-90%) in humans (21–23). Additional orthornaviruses in the families *Paramyxoviridae* (mumps, measles and parainfluenza viruses) (24–26), and *Flaviviridae* (yellow fever, Dengue and Zika viruses) (27), are also major contributors to human disease.

Since the work of Katzourakis and Gifford in 2010, the diversity of vertebrate EVEs at the level of multiple viral families has not been systematically surveyed (1). We took advantage of the larger sequence data sets available today together with a cloud-computing approach, to carry out a comprehensive search for non-retroviral EVEs (kingdoms *Shotokuvirae* and *Orthornavirae*) in vertebrate genomes. Using 24,478 viral protein queries, we identified 2,040 EVEs in 295 host species. These include the first EVEs belonging to the families *Nairoviridae*, *Paramyxoviridae*, *Chuviridae* and *Benyviridae* in vertebrate genomes, and from the *Hepacivirus* genus of flaviviruses. We also discovered endogenous ectodomains closely related to those found in reptarenaviruses and filoviruses, which suggest a macroevolutionary scenario for the origin of glycoprotein ectodomains. Our analysis sheds light on the evolutionary history and ecology of multiple viral lineages, and shows the value of cloud-computing for revealing the diversity of EVEs in vertebrate genomes.

## Results

We identified a total of 2,040 EVEs in the genome assemblies of 295 vertebrates, in addition to 17 exogenous virus sequences (Supplementary figures 1 and 2, Supplementary excel file 1). Among these sequences, we report the first non-retroviral EVEs in vertebrate genomes belonging to the families *Chuviridae* (121 EVEs), *Paramyxoviridae* (19 EVEs), *Benyviridae* (22 EVEs) and *Nairoviridae* (1 EVE). We found the first evidence of an EVE from the *Hepacivirus* genus of flaviviruses (initially 4 EVEs, extended to 21 EVEs). We also identified close hits to the ectodomains of reptarenaviruses in tarsier genomes, and to the ectodomains of filoviruses in the genomes of cartilaginous fish and the Komodo dragon, contained within retrovirus-like elements.

### Chuvirus EVEs in the genomes of fish, mammals and non-avian reptiles

Chuviruses are negative-sense RNA viruses (Order *Jingchuvirales*) described mainly from metagenomic samples (28). They have been found in arthropods and associated with a number of vertebrates (28). Although chuvirus-like EVEs had been described in a number of arthropod genomes (29,30), the nature of the vertebrate associated viruses remained unclear. We found 28 EVEs similar to the RNA-dependent RNA polymerase in teleost fish, and 92 EVEs similar to the nucleoprotein in teleosts, amphibians, snakes and lizards (lepidosaurs), and marsupials (Figure 1). The vertebrate-associated chuviruses form a well-supported clade with the chuvirus EVEs (posterior probability = 1) in the RdRp phylogeny (Figure 1A), and occupy central nodes in the nucleoprotein network surrounded by the chuvirus EVEs found in vertebrates (Figure 1B). Examination of EVE loci from teleosts and marsupials revealed that some of these integrations are orthologous and date back to 11.9 – 35 million years ago (MYA).

**Figure 1.**
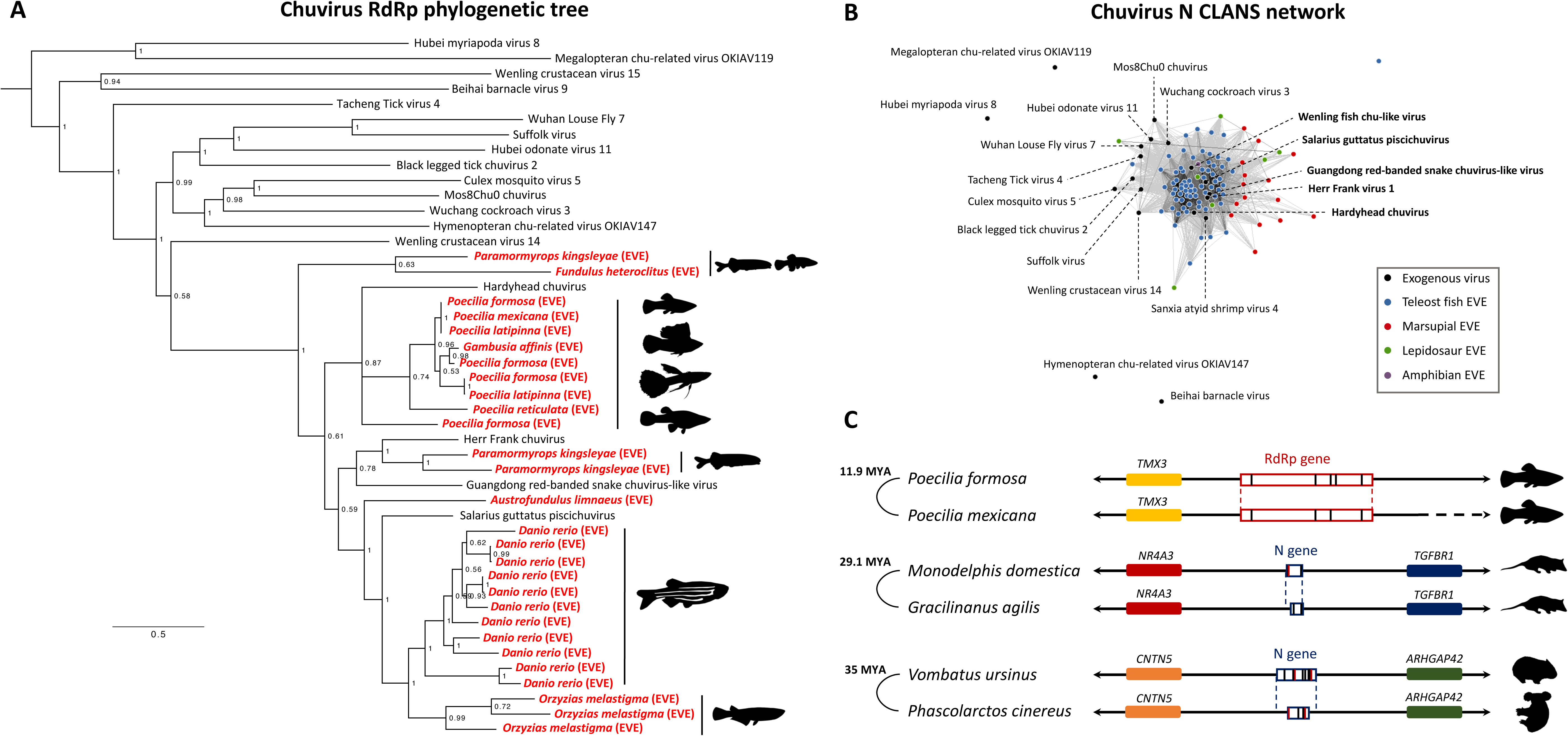
Chuvirus EVEs in vertebrate genomes. (**A**) Bayesian phylogenetic tree of the RdRps of exogenous chuviruses and the EVEs found in teleost fish (in red). Some species have multiple integrations suggesting a close interaction with these viruses. Note how the vertebrate-associated viruses and EVEs form a clade that is paraphyletic to the chuviruses found in arthropods. The tree was rooted with Hubei myriapoda virus 8 (*Myriaviridae*) and Megalopteran chu-related virus 119 (*Crepuscoviridae*) as outgroups. Tree inferred in MrBayes3 using the LG+F+I+G4 model and 4.74M generations (relative burn-in = 25%). EVEs are shown in red. **(B)** CLANS network of the nucleoprotein of exogenous chuviruses, vertebrate chuvirus EVEs and the two outgroups mentioned above. Edges are drawn between nodes with a significance of p < 1e-15. The vertebrate EVEs are well connected to the central network that includes vertebrate-associated chuviruses and a number of chuviruses from arthropods. **(C)** Syntenic arrangement of the most proximal genes was used to establish orthology of three integrations. Vertical red bars in the EVEs indicate internal stop codons, while black bars indicate indels. The minimum date of integrations in each species pair is based on the divergence of the host species in TimeTree (31).

### Paramyxovirus EVEs in the genomes of teleost fish

Paramyxoviruses are nonsegmented, negative-sense RNA viruses classified in the order *Mononegavirales* (32). Although paramyxoviruses infect a wide variety of vertebrate hosts (32), EVEs from paramyxoviruses had not been described. We found 17 EVEs similar to the RdRp of paramyxoviruses, and 2 EVEs similar to the nucleoprotein in the genomes of teleost fish. Multiple integrations were found in species of fish from the family Labridae (*Labrus*, *Notolabrus*, *Cheilinus*), in the leopard coral grouper *Plectropomus leopardus* (Serranidae), and in the Mexican tetra *Astyanax mexicanus* (Characidae). Phylogenetic analysis placed most of the RdRp EVEs in a clade with Wenzhou pacific spadenose shark paramyxovirus (posterior probability = 0.9), while a single EVE from the coral grouper was placed between this clade and a clade composed of more widely known paramyxoviruses such as Measles virus, Hendra virus or human respiroviruses (Figure 2A). Structural comparison of an open reading frame fragment found in the genome of the Mexican tetra to *Orthorubulavirus mammalis*, revealed a conserved structure of the RdRp (Figure 2B). Interestingly, the nucleoprotein-like EVEs found in the Mexican tetra were placed next to a group of bat paramyxoviruses (Figure 2C).

**Figure 2.**
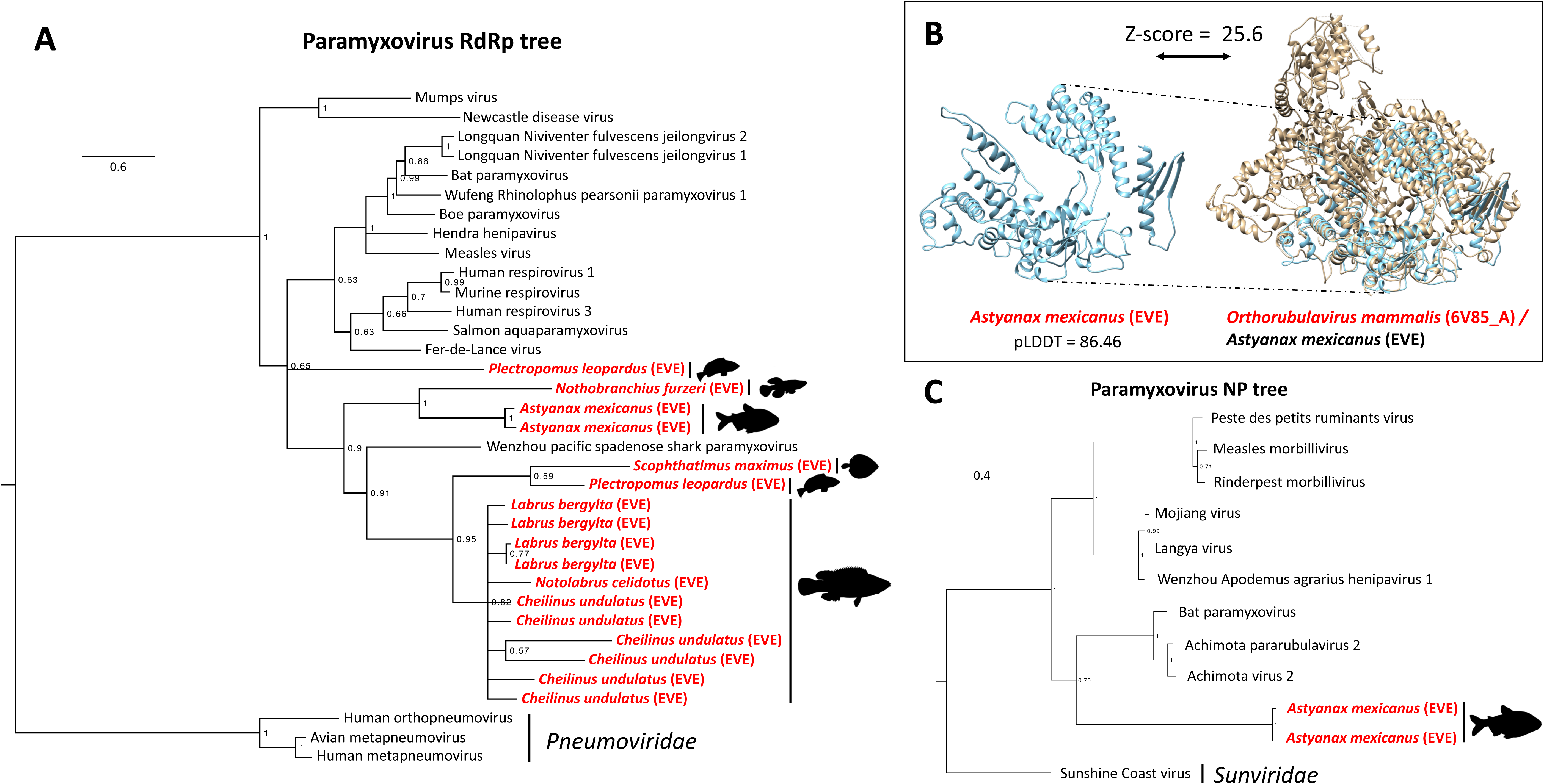
Paramyxovirus EVEs in the genomes of teleost fish. (**A**) Bayesian tree of the RdRp of exogenous paramyxoviruses and the EVEs found in teleost fish (in red). Most EVEs form a clade together with Wenzhou pacific spadenose shard paramyxovirus. The tree was outgroup-rooted with RdRp sequences from pneumoviruses. Tree inferred in MrBayes3 using the LG+F+I+G4 model and 9.06M generations (relative burn-in = 25%). EVEs are shown in red. **(B)** Predicted structure of an RdRp fragment present in the genome of the Mexican tetra and comparison to the RdRp structure of Parainfluenza virus 5 (*Orthorubulavirus mammalis*). **(C)** Bayesian tree of the nucleoprotein of paramyxoviruses and the EVEs found in the Mexican tetra. The EVEs are nested within the *Paramyxoviridae* with high support (posterior probability = 1), and are closest to a group of bat paramyxoviruses with a posterior probability = 0.75. Tree inferred in MrBayes3 using the LG+I+G4 model and 1M generations (relative burn-in = 25%). EVEs are shown in red.

### Plant and fungal-like EVEs (family *Benyviridae*) in vertebrate genomes

Benyviruses are multipartite, positive-sense RNA viruses which have been known to infect plants (33), but more recently have also been isolated from fungi and some insects (34). We found 19 EVEs with similarity to the RdRp of benyviruses in the genomes of caecilians (*Rhinatrema*, *Microcaecilia*), lizards (*Podarcis*, *Gekko*), snakes (*Python*), the West African lungfish (*Protopterus annectens*) and the Great white shark (*Carcharodon carcharias*). In the phylogeny of benyvirus RdRps (Figure 3), the EVEs of vertebrates were placed in a clade with two benyviruses isolated from insects (Diabrotica undecimpunctata virus 2, and Bemisia tabaci beny-like virus 6), thus forming a clade of animal viruses. The phylogeny also recovered a clade of benyviruses that infect land plants and another that infects mostly fungi (with the exception of some viruses isolated from the silverleaf whitefly, *Bemisia tabaci*). A tanglegram of the benyvirus RdRps and the host phylogeny, was able to recover the split between land plants and fungi + animals (Opisthokonta). In the animal infecting group, the inconsistency of both phylogenies suggest a dynamic history of cross-species transmissions (Figure 3). We also found 6 EVEs with similarity to the coat protein of benyviruses in lizards (*Podarcis*, *Lacerta*, *Zootoca*), and the small-spotted catshark (*Scyliorhinus canicula*).

**Figure 3.**
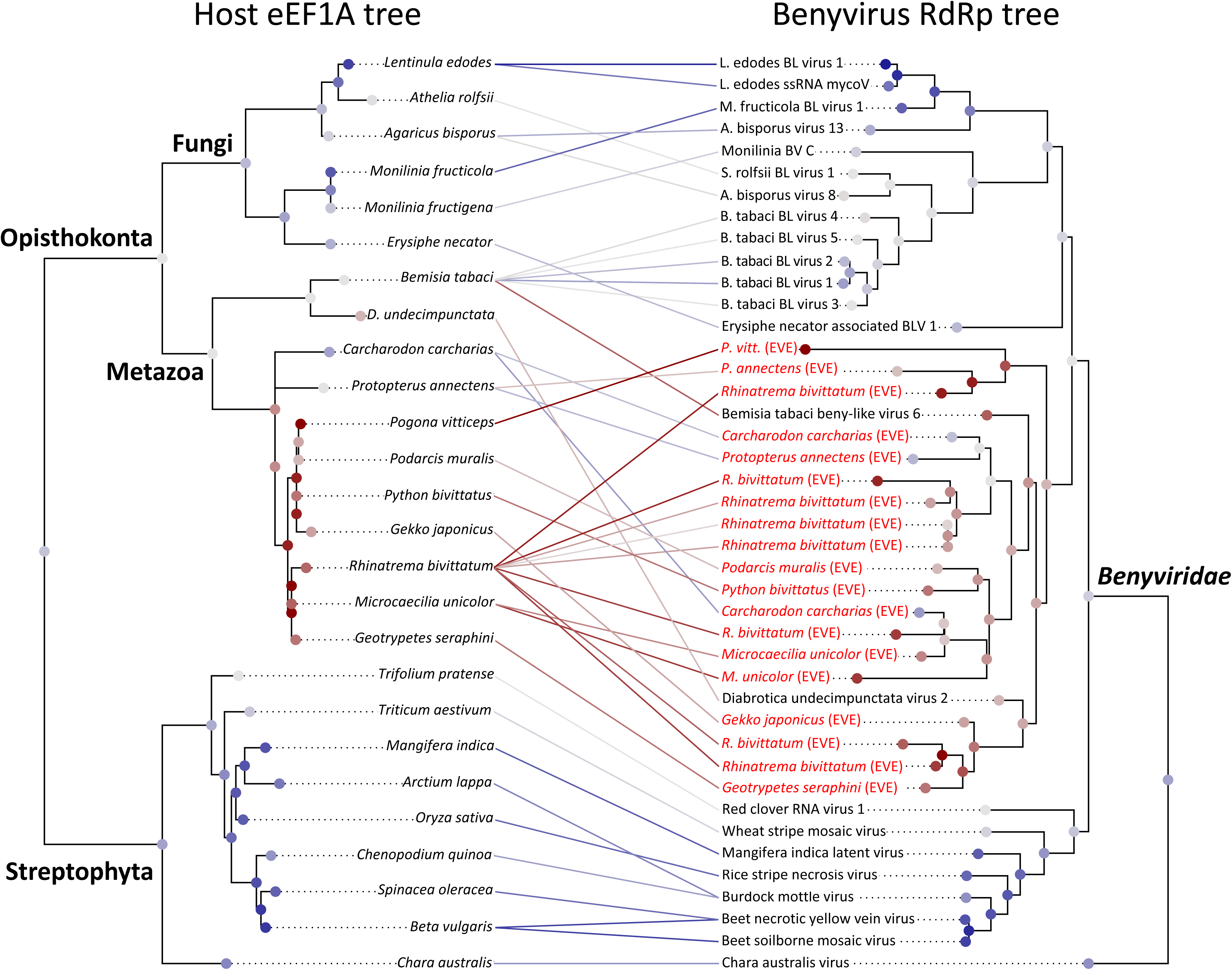
Tanglegram of the phylogenies benyviruses (including vertebrate EVEs) and their eukaryotic hosts. The benyvirus RdRp and host phylogenies point at deep codivergences and more recent cross-species transmissions in the three main groups (plant, fungi, animal benyviruses). The more basal position of Chara australis virus in the RdRp phylogeny could be interpreted as an ancient virus jump between photosynthetic organisms and the ancestors of animals and fungi (Opisthokonta). The maximum-likelihood trees were inferred in RAxML-NG (eEF1A: LG+I+G4, RdRp: LG+F+I+G4), and the tanglegram inferred using the maximum incongruence algorithm (MIC) in RTapas. EVEs are shown in red.

### Nairovirus EVE in the genome of the Etruscan shrew

Nairoviruses are negative-sense RNA viruses with 3 genomic segments S, M and L. The S segment carries the gene that encodes the nucleoprotein (35). Nairoviruses infect arthropods and can be transmitted to humans via tick bites (35). Some nairoviruses can cause disease in humans, but the Crimean-Congo Haemorrhagic Fever (CCHF) viruses are noteworthy for being highly pathogenic (36). Previously, EVEs similar to the nucleoprotein of nairoviruses had been described in the genome of the black-legged tick *Ixodes scapularis* (1). However, they were distantly related to the nucleoproteins of CCHF viruses. We found an EVE in the genome of the Etruscan shrew (*Suncus etruscus*), which is the closest EVE to CCHF virus and which can be placed in the same genus, *Orthonairovirus* (Figure 4A). Using this sequence to query the nr protein database (NCBI), we were able to identify new EVEs in the genomes of additional species of ticks (*Rhipicephalus sanguineus*, *Dermacentor silvarum*, *Dermacentor andersoni*), and in other chelicerates (scorpions and spiders). Comparison of the predicted EVE protein structures, show the high similarity between the nucleoproteins from the Etruscan shrew EVE and CCHFV, and between the black-legged tick and South Bay virus (Figure 4B).

**Figure 4.**
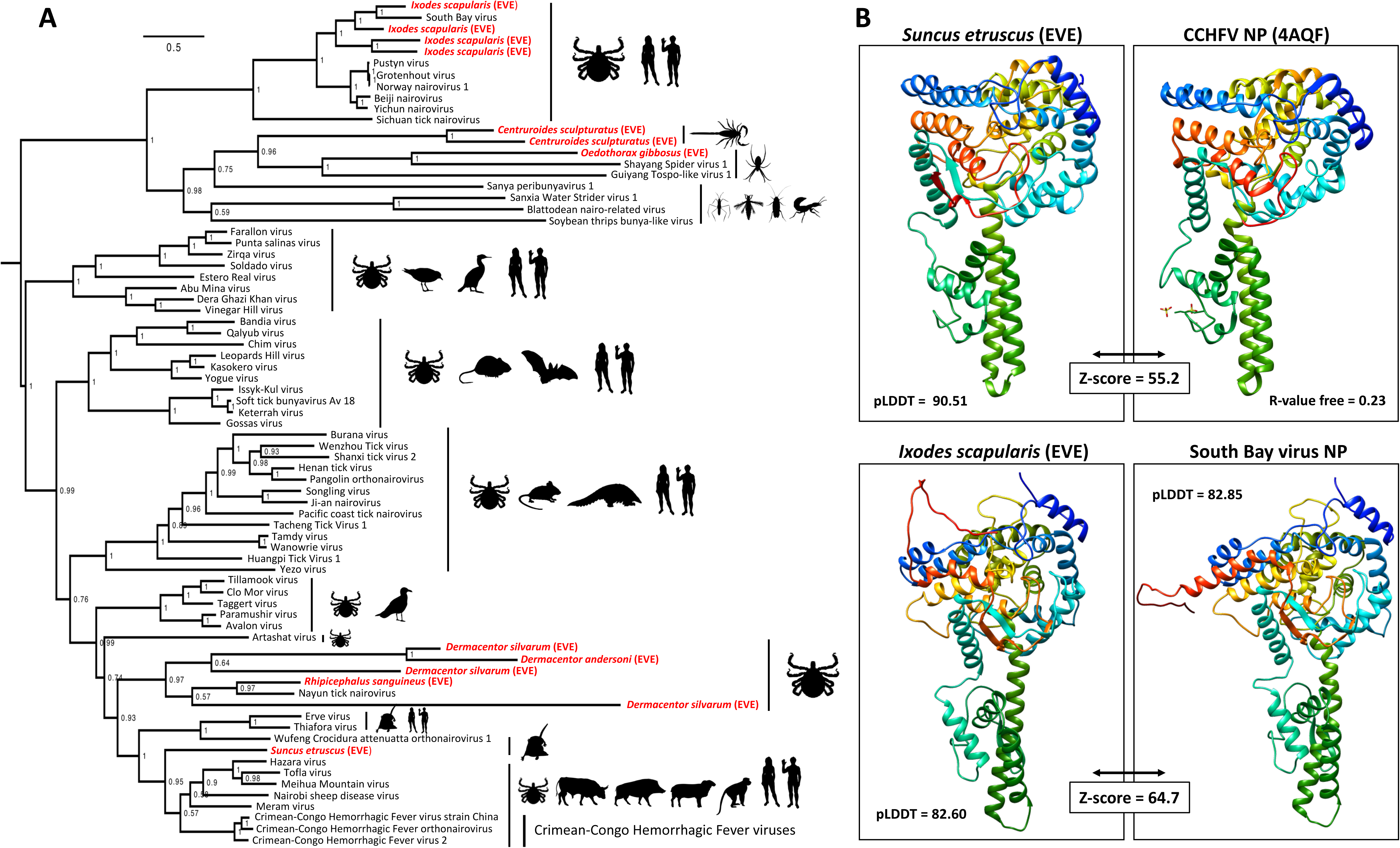
Nairovirus EVEs in the genome of the Etrsucan shrew and ticks. (**A**) Bayesian phylogeny of the nairovirus nucleoprotein gene including EVEs from the Etrsucan shrew, ticks and other chelicerates, together with exogenous nairoviruses. The element found in the Etrsucan shrew genome forms a clade with the Crimean Congo Hemorrhagic Fever viruses/Haza virus, sister to the Erve/Thiafora and Wufeng Crocidura attenuatta orthonairovirus 1 clade, known to infect soricid shrews of the subfamily *Crocidurinae*. Tree inferred in MrBayes3 with a codon-partitioned model (1^st^ and 3^rd^ positions: GTR+G4, 2^nd^ position: GTR+I+G4), and 5M generations (relative burn-in = 25%). EVEs are shown in red. **(B)** Structural comparison of nucleoproteins from EVEs in the Etrsucan shrew and black-legged tick genomes with exogenous nairoviruses. Structures were modelled in Alphafold2 to a good backbone accuracy (pLDDT > 80) or downloaded from PDB. The Etruscan shrew element adopts a structure highly similar to the structure of Crimean-Congo Hemorrhagic Fever virus determined by X-ray crystallography. The black-legged tick predicted structure is more similar to the South Bay virus structure as predicted from phylogenetic analysis.

### *Hepacivirus* EVE in the genomes of murine rodents

Hepaciviruses are positive-sense RNA viruses in the family Flaviviridae, which are classified in the genus Hepacivirus (37). People chronically infected with Hepatitis C virus (HCV) are at a significant risk of liver disease including fibrosis, cirrhosis and hepatocellular carcinoma (38). We found hits homologous to a ∼67 aa fragment of the positive-sense single-stranded RNA (ps-ssRNA) polymerase domain (Superfamily cl40470) of Rodent hepacivirus ETH674/ETH/2012, in the genomes of rodents in the subfamily Murinae (Figure 5A, 5B). Examination of the genomic context across 21 species, showed that the integration was orthologous but degraded in murine genomes (Figure 5C). Given that the hepacivirus EVE is shared between mice (*Mus* spp.) and rats (*Rattus* spp.), this suggests a minimum age of 11.7–14.2 MYA. Intriguingly, we have only been able to identify this sequence in the polymerase domain of the Rodent hepacivirus ETH674/ETH/2012, isolated from the Ethiopian white-footed mouse (*Stenocephalemys albipes*).

**Figure 5.**
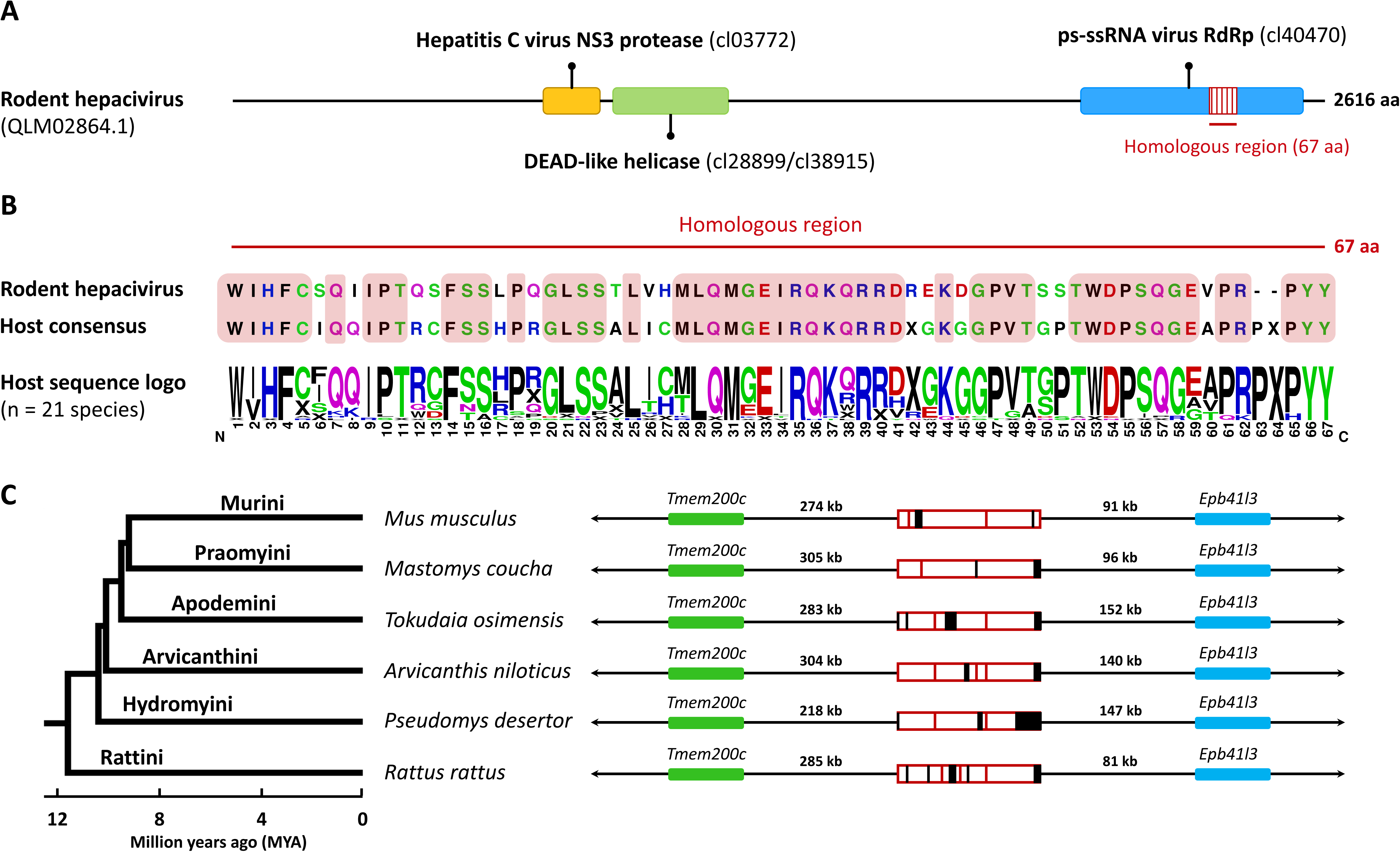
Hepacivirus EVE in the genomes of rodents from the subfamily *Murinae*. (**A**) Conserved domain annotation of the Rodent hepacivirus ETH674/ETH/2012 (QLM02864.1) polyprotein. The region of homology to the EVEs is embedded within the ps-ssRNA domain. **(B)** Comparison of the region of homology between Rodent hepacivirus ETH674/ETH/2012 and the consensus sequence obtained from 21 murine genomes. Identical amino acids at a given position are highlighted in a red box (the two sequences are 75% pair-wise identical at the amino acid level). The sequence logo shows variation at the given position proportional to frequency (0-100%). **(C)** Orthology across 5 representative species in 5 tribes (Murini, Praomyini, Apodemini, Arvicanthini, Hydromyini, Rattini) of the subfamily Murinae, together with a phylogeny of the group. Flanking genes were identified in the mouse (*Mus musculus*) assembly, and used to annotate the region in the other assemblies. Red bars: internal stop codons, black rectangles: indel mutations.

### Ancient captures of the retroviral ectodomain by filoviruses and reptarenaviruses

The envelope proteins of retroviruses, and the glycoproteins of some filoviruses (Ebolavirus, Marburgvirus, Cuevavirus, Dianlovirus and Tapjovirus), contain an ectodomain with heptad-repeat sequences and an immunosuppressive domain (ISD) region (39). Interestingly, the glycoproteins of arenaviruses in the genus Reptarenavirus also contain a similar ectodomain (40). We found hits closely related to the ectodomain of reptarenaviruses in the genomes of the Philippine tarsier (*Carlito syrichta*) and the Western Tarsier (*Cephalopachus bancanus*) (Supplementary excel file 1). We noticed that these hits were in close proximity to other retroviral domains (gag, RT, RNaseH, rve), they were flanked by direct repeats, and occurred at the expected relative position of the *env* gene, establishing that these were hits to retroviral elements. By searching for other hits related to filovirus and reptarenavirus ectodomains, we found additional ectodomains surrounded by retroviral features (or annotated as such) in the genomes of lizards (*Mabuya*, *Varanus*), and cartilaginous fish (*Chiloscyllium*, *Scyliorhinus*, *Amblyraja*, *Leucoraja*). After confirming that additional retrovirus ectodomains fell outside this clade, we focused on the ingroup to construct a time-calibrated tree (Figure 6, Supplementary figure 4).

**Figure 6.**
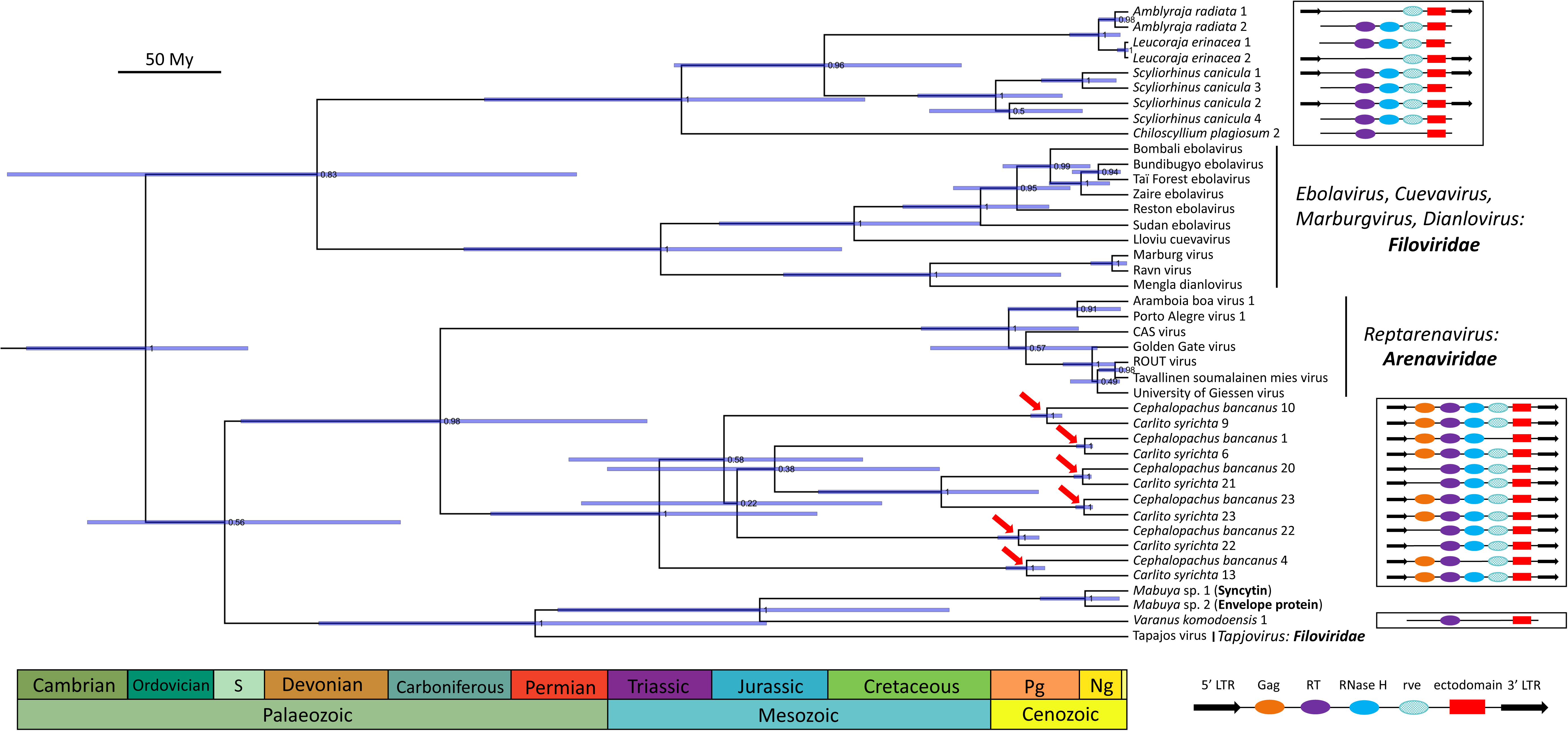
Bayesian timetree of the ectodomain homologues found in retroviruses, filoviruses and reptarenaviruses. The ectodomains of reptarenaviruses form a highly supported clade (posterior probability = 0.98) with the endogenous ectodomains found in tarsiers (*Carlito syrichta*, *Cephalopachus bancanus*). The ectodomains of ebolaviruses, cuevaviruses, marburgviruses and dianloviruses, form a clade which is the sister group to the endogenous ectodomains found in cartilaginous fish. However, the ectodomain of Tapajos virus forms a distinct clade (posterior probability = 1) with endogenous ectodomains found in lizards (*Mabuya*, *Varanus*), suggesting that the Tapajos virus ectodomain was captured independently from the ectodomains of other filoviruses. The tree was inferred in BEAST2 with the JTT+G4 site model, using the Optimised Relaxed Clock (ORC) and 20M generations (relative burn-in = 25%). The red arrows indicate pairs of tarsier orthologues. A diagram with the genomic context of the endogenous ectodomains is shown to the right, and suggests that the endogenous ectodomains form part of endogenous retroviral elements.

The posterior evolutionary rate of the ectodomains was estimated at 3.2 × 10^-9^ amino acid substitutions per site per year (± 4.4 x 10^-10^ aa subs./site/year, Supplementary figure 5). This is consistent with the higher neutral evolutionary rates reported for immunoglobulin kappa (3.7 × 10^-9^ aa subs./site/year) and gamma C chains (3.1 × 10^-9^ aa subs./site/year), and the complement C3a anaphylatoxin (2.7 × 10^-9^ aa subs./site/year) (41). It is also consistent with the time-dependency of viral evolutionary rates, which tend to converge on the host rate over geological timescales (42). These observations indicate that the timescale of evolution was calibrated properly; misspecified priors would have resulted in a significant departure from the time-dependent and neutral expectations.

In the Bayesian phylogeny (Figure 6), the ectodomains of reptarenaviruses were placed as the sister group to the ectodomains in tarsiers with high confidence (posterior probability = 0.98). The ectodomains from ebola-, cueva-, marburg– and dianloviruses were placed as the sister clade to the ectodomains of retroelements found in cartilaginous fish (posterior probability = 0.83). On the other hand, the ectodomain from the filovirus Tapajos virus (*Tapjovirus*), which was found in the venom gland of the Common lancehead viper (*Bothrops atrox*) (43), was placed forming a strongly supported clade with ectodomains found in lizard retroelements (posterior probability = 1). These findings suggest that ectodomains have been captured from retroviral elements 3 times independently, twice by filoviruses and once by reptarenaviruses, over a timescale of hundreds of millions of years.

## Discussion

We discovered novel EVEs in vertebrate genomes belonging to the families *Chuviridae*, *Paramyxoviridae*, *Benyviridae* and *Nairoviridae*. This represents a 44% increase in the family-level diversity of vertebrate non-retroviral EVEs (9 to 13 families). In addition, we identified the first *Hepacivirus* EVE in the genomes of murine rodents, and found retroviral elements with ectodomains related to those of reptarenaviruses and filoviruses. Endogenous viral elements in the families *Circoviridae*, *Parvoviridae*, *Bornaviridae, Filoviridae* and *Flaviviridae*, accounted for 91% of the EVEs (1,858/2,040) found during our search. Therefore, in a single systematic search, our strategy allowed for both increased sensitivity and detection of novel and less abundant EVEs (9%), as well as reproduction of previous and recent findings in the field.

Chuviruses, which are a family of RNA viruses discovered in metagenomes, have been found associated mainly with arthropods (28). Chuvirus EVEs have been described in the genomes of arthropods, further supporting infection of this group of invertebrates by chuviruses (30). A number of chuviruses have also been found associated with vertebrates, but having been isolated only from metagenomic samples, the nature of the association with vertebrates was uncertain (44). We show evidence that chuviruses actively infect vertebrates, by the discovery of 121 EVEs in teleost fish, lepidosaurs, amphibians and marsupials. The vertebrate-associated chuviruses formed a clade with the chuvirus EVEs in vertebrates (posterior probability = 1), strongly supporting that there is a vertebrate-specific clade of chuviruses. The detection of orthology of several chuvirus EVEs on the order of 11-35 million years, indicate that chuviruses have infected vertebrates from at least the Eocene epoch. These results are in line with recent evidence that chuviruses can infect and cause lymphocytic meningoencephalomyelitis in wild species of turtles (44).

Surprisingly, we found 22 vertebrate EVEs that could be assigned to the family *Benyviridae*. Benyviruses are plant pathogens, but a few viruses have been identified from insect metagenomes (45,46). Our study uncovered endogenous benyviruses in vertebrate genomes, which form an animal-specific clade with four benyviruses isolated from insects (*Diabrotica undecimpunctata*, *Sesamia inferens* and *Harmonia axyridis*, Supplementary figure 3). This implies that a clade of benyviruses exhibits tropism for animals, extending its range to a new host kingdom. As shown in Figure 3, the benyviruses of animals seem to undergo frequent cross-species transmissions. Additionally, we uncovered 19 EVEs from paramyxoviruses in both freshwater and marine teleost fish. Paramyxoviruses are known to infect fish (47), and some have been associated with disease including epidermal/gill necrosis, gill inflammation and buccal/opercular haemorrhage (48). Our results highlight the need to better characterise the diversity of paramyxoviruses in fish hosts, in particular pointing to close interactions with the orders Perciformes (most diverse order of fish), Cyprinodontiformes (toothcarps) and Pleuronectiformes (flatfish).

We provide the first evidence for an EVE from the genus *Hepacivirus* in murine rodents. This EVE shares high homology (75% amino acid identity) across a segment of the polymerase domain with Rodent hepacivirus ETH674/ETH/2012. Further confirmation of orthology across rodents of the Murinae subfamily, constitute direct evidence that hepaciviruses have infected murine rodents for at least 11.7–14.2 million years. Rodents in the subfamily Murinae are inferred to have shared a most recent common ancestor in Southeast Asia 15.9 (14.1–18.2) MYA (49), while the sequence of Rodent hepacivirus ETH674/ETH/2012 was isolated from an Ethiopian white-footed mouse (*Stenocephalemys albipes*) in Africa (50), suggesting a close coevolutionary history with murine rodents. These observations agree with recent findings that highlight murid rodents as important hepacivirus hosts (50,51), together with molecular estimates based on present-day sequences that suggest an origin of the *Hepacivirus* genus ∼22 million years ago (51). Given that the homologous sequence found in Rodent hepacivirus ETH674/ETH/2012, and the murine rodent EVE seem to be a unique derived feature (synapomorphy), it appears likely that hepaciviruses as a whole are older than 22 million years, which can be considered a minimum conservative estimate.

Although nairovirus-like EVEs had been described in black-legged ticks (*Ixodes scapularis*) (1), we identified the first vertebrate nairovirus EVE in the genome of the Etruscan shrew (*Suncus etruscus*). Discovery of this element points to the importance of shrews as reservoirs of potentially pathogenic orthonairoviruses. This EVE is the closest to the group of the Crimean-Congo Hemorrhagic Fever (CCHF) viruses, and sits between this clade and a group which includes Erve virus which is suspected to cause severe headache and intracerebral haemorrhage in humans (52). The related Erve and Thiafora viruses found in France and Senegal, were initially isolated from shrews (*Crocidura russula*, *Crocidura* sp.) (53,54). A number of recently discovered orthonairoviruses have also been isolated from shrews including: Wufeng orthonairovirus 1 from *Crocidura attenuata* in China, Lamusara and Lamgora viruses from *Crocidura goliath* in Gabon (55), and Cencurut virus from *Suncus murinus* in Singapore (56). These data indicate that shrews in the subfamily Crocidurinae are important natural reservoirs of orthonairoviruses in Europe, Africa and Asia. Similarly, our discovery of EVEs related to Nayun tick nairovirus in *Rhipicephalus sanguineus*, *Dermacentor andersoni* and *Dermacentor silvarum*, implicate these tick species as additional vectors of orthonairoviruses. This agrees with the isolation of Nayun tick nairovirus from a *Rhipicephalus* tick (57). Together, these observations suggest a close interaction between multiple species of ticks with nairoviruses, and support the role of crocidurine shrews as important mammalian reservoirs for orthonairoviruses.

There is potential for non-retroviral EVEs to function in EVE-derived immunity. In the thirteen-lined squirrel (*Ictidomys tridecemlineatus*), an endogenous bornavirus-like N gene (416 aa long) can inhibit Borna disease virus (BDV) replication, and block *de novo* infection by BDV (58). Recently, a parvoviral-like Rep gene in the genome of degus (*Octodon degus*), encoding a 508 amino acid product, was shown to inhibit replication of the model parvovirus Minute virus of mice (MVM) (59). We noticed that multiple EVEs in our data set contain large (>400 amino acid) open reading frames, which show similarity to nucleoprotein and polymerase genes of exogenous viruses. In particular, we describe EVE loci for the families *Nairoviridae*, *Paramyxoviridae* and *Chuviridae*, which seem like interesting candidates for exploration of potential EDI function. However, some of these genes may have acquired other unexpected functions in host biology. For example, in pea aphids (*Acyrthosiphon pisum*), expression of an endogenous densovirus (*Parvoviridae*) EVE has been co-opted to trigger wing development as an environmentally plastic trait (60).

Our findings also shed light on the origin of ectodomains in the glycoproteins of filoviruses and reptarenaviruses. The presence of an ectodomain containing an immunosuppressive region in Ebola and Marburg viruses, and homology to the ectodomain of retroviruses, had been noted by Bénit et al. (39). Similarly, the glycoproteins of reptile arenaviruses (genus *Reptarenavirus*), were reported to be highly similar to the glycoproteins of filoviruses (40). We could not detect presence of the ectodomain in fish filoviruses (*Oblavirus*, *Striavirus*, *Thamnovirus*), nor in other arenaviruses aside from *Reptarenavirus*. This patchy distribution suggests that presence of the ectodomain is a derived character (apomorphy) in some filoviruses and *Reptarenavirus*, and not an ancestral trait for the families *Filoviridae* and *Arenaviridae*. Here, we propose a macroevolutionary scenario whereby retroviral ectodomains were captured by filoviruses and arenaviruses three times independently: 1) by the common ancestor of *Ebolavirus*, *Marburgvirus*, *Cuevavirus* and *Dianlovirus*, 2) by Tapajos virus (or its direct ancestor), and 3) by the common ancestor of reptarenaviruses. This degree of convergence argues in favour of a strong selective advantage gained by acquisition of the ectodomain, probably driven by improved suppression of the tetrapod immune system.

Our study demonstrated the capacity of cloud-based, highly parallelised approaches to harness the vast amounts of sequence data, revealing novel insights into the biology of viruses. Specifically, we increased the diversity of non-retroviral EVEs known in vertebrate genomes from 9 to 13 families, and presented the first evidence of endogenous chuviruses, paramyxoviruses, plant-like viruses (benyviruses), orthonairovirus and hepacivirus in vertebrate genomes. These results suggest the extension of the host range of chuviruses and benyviruses to vertebrates, and highlight the close evolutionary association of crocidurine shrews and murine rodents with orthonairoviruses and hepaciviruses, respectively. We also propose a macroevolutionary model for the acquisition of ectodomains in filovirus and reptarenavirus glycoproteins from a retroviral source. These discoveries open rich grounds to study the potential function of diverse non-retroviral EVEs on host biology. We foresee that with ever increasing availability in genomic sequence data, and the advance in computing power and algorithms, our knowledge of the genomic fossil record of viruses will continue to increase.

## Methods

We used cloud computing on the Google Cloud Platform, to search for homology to a comprehensive set of protein sequences derived from viruses in the kingdoms *Shotokuvirae* (ssDNA and dsDNA viruses) and *Orthornavirae* (RdRp-containing RNA viruses), across all representative vertebrate genomes. Hits were extracted and processed for taxonomic assignment into their respective viral groups (hits that did not return 50% reciprocal hits to viruses were considered ambiguous and not considered further). Hits showing high sequence similarity to known viruses or otherwise present in small contigs (<10,000 bp) without nearby host genes were considered exogenous viruses. Confirmed endogenous viral elements were then annotated, aligned and used in phylogenetic inference together with homologues from exogenous viruses. A more detailed description of the methods is described in the following sections.

### Selection of viral queries and sequence clustering

We downloaded 439,594 protein sequences from complete viral genomes available at NCBI Virus (61) during September, 2022. The sequences were partitioned according to their viral family and clustered using MMSeqs2 (62). Clustering was performed using a minimum pairwise identity (--min_seq_id) of 65% at the amino acid level and the default cover (80%). Sequence centroids were extracted from each cluster and used as representative sequences for downstream analyses. This representative set contained 24,478 sequences.

### Elastic-BLAST searches on the Google Cloud Platform

Cloud searches for each viral family were conducted on the Google Cloud Platform (63) using the Elastic-BLAST algorithm (64) in September, 2022. Each search was performed with tblastn (tblastn-fast option) against the entire database of representative vertebrate genomes (ref_euk_rep_genomes, taxid: ‘7742’), and using an e-value of 1e-5. The output was saved in tabular format (-outfmt ‘7’). The analysis returned 196,899 hits to the viral queries.

### Curation of non-redundant loci

Hits to host genomes were merged with bedtools2 (65) in order to reduce redundancy in the data set. Strictly overlapping hits and hits that were at a maximum distance of 200 nt (based on their genomic coordinates) were merged to give a single range in the host genome (-d 200). We thus obtained a set of 26,324 non-redundant genomic regions. We then downloaded the genomic sequences from the merged ranges in fasta format using efetch (66).

### DIAMOND reciprocal searches and taxonomic assignment

To assess the origin of the host sequences (whether viral or host), we downloaded and compiled the complete non-redundant (nr) protein database with taxonomic information on the High-Performance Computing cluster at the University of Oxford. We then performed a reciprocal similarity search using the host sequences as queries and the nr database with DIAMOND blastx (67), keeping only the top 25 hits. We obtained 558,589 reciprocal hits in total. Next, we used custom scripts written in Python 3 to parse the taxonomic labels obtained for each query sequence and assign them to the majority-rule viral family. Sequences were considered viral if ≥50% of the reciprocal hits were to “Viruses”. Viral sequences falling on short contigs or with high similarity to known exogenous viruses (>99% identical) were considered exogenous viruses present in the assemblies (and not EVEs).

### Phylogenetic inference and structural predictions

We focused on elements which had not been described as EVEs in the literature for the phylogenetic and structural analyses. Predicted protein sequences for each locus were obtained and annotated manually using blastx / conserved domain search on the NCBI web server (68–70), GeneWise on the EBI web server (71,72) or HHpred on the Max Planck Institute’s web server (73,74). Exogenous virus homologues were searched against the nr database using blastp online. Multiple sequence alignments were obtained using MAFFT (75) or MACSE (76). Trees were estimated from amino acid data, except for nairoviruses which were based on a nucleotide alignment. We selected the best substitution models in Modeltest-NG (77). Trees were estimated in RAxML-NG (78) with 200 starting trees and up to 2,000 bootstraps (autoMRE{2000}), until convergence in MrBayes3 (79) (standard deviation of split frequencies < 0.01) and in BEAST2 (80) (after inspecting the runs for good mixing, stationarity and effective sample sizes > 200). For the inference of the time tree of ectodomains, we used orthology of the tarsier elements and their estimated ages (based on LTR divergence, Supplementary excel file 2) to calibrate internal nodes in the tree, and used a prior distribution on the root of the tree assuming that the retroelements present in cartilaginous fish/tetrapods codiverged with their gnathostome hosts (prior mean 462, prior 95% CI: 436-489 MYA). Cophylogenetic analysis for benyviruses was performed and plotted in RTapas using the maximum incongruence algorithm (81). We predicted select paramyxovirus and nairovirus protein structures for *de novo* using AlphaFold2 (82) as implemented in ColabFold (83). We used amber relaxation on the top ranked structure, and either 24 or 48 recycles. Network analysis of chuvirus capsid proteins was performed using CLANS 2.0 (84,85), with a p-value < 1e-15.

## Acknowledgements

A.K. discloses support for the research and publication of this work from the European Research Council (grant number 101001623-PALVIREVOL). J.G.N.B. acknowledges support in computing credits and access to resources from Google Cloud (grant number EDU Credit 212888085). The authors would also like to acknowledge the use of the University of Oxford Advanced Research Computing (ARC) facility in carrying out this work. http://dx.doi.org/10.5281/zenodo.22558.

## Supplementary materials

All data and code supporting this work are available at the Open Science Framework server: https://osf.io/7rqa2/.

## Notes

https://osf.io/7rqa2/

### Competing Interest Statement

The authors have declared no competing interest.

