## Supplementary material for "Deep-mining of vertebrate genomes reveals an unexpected diversity of endogenous viral elements": Supplementary_figures.pdf

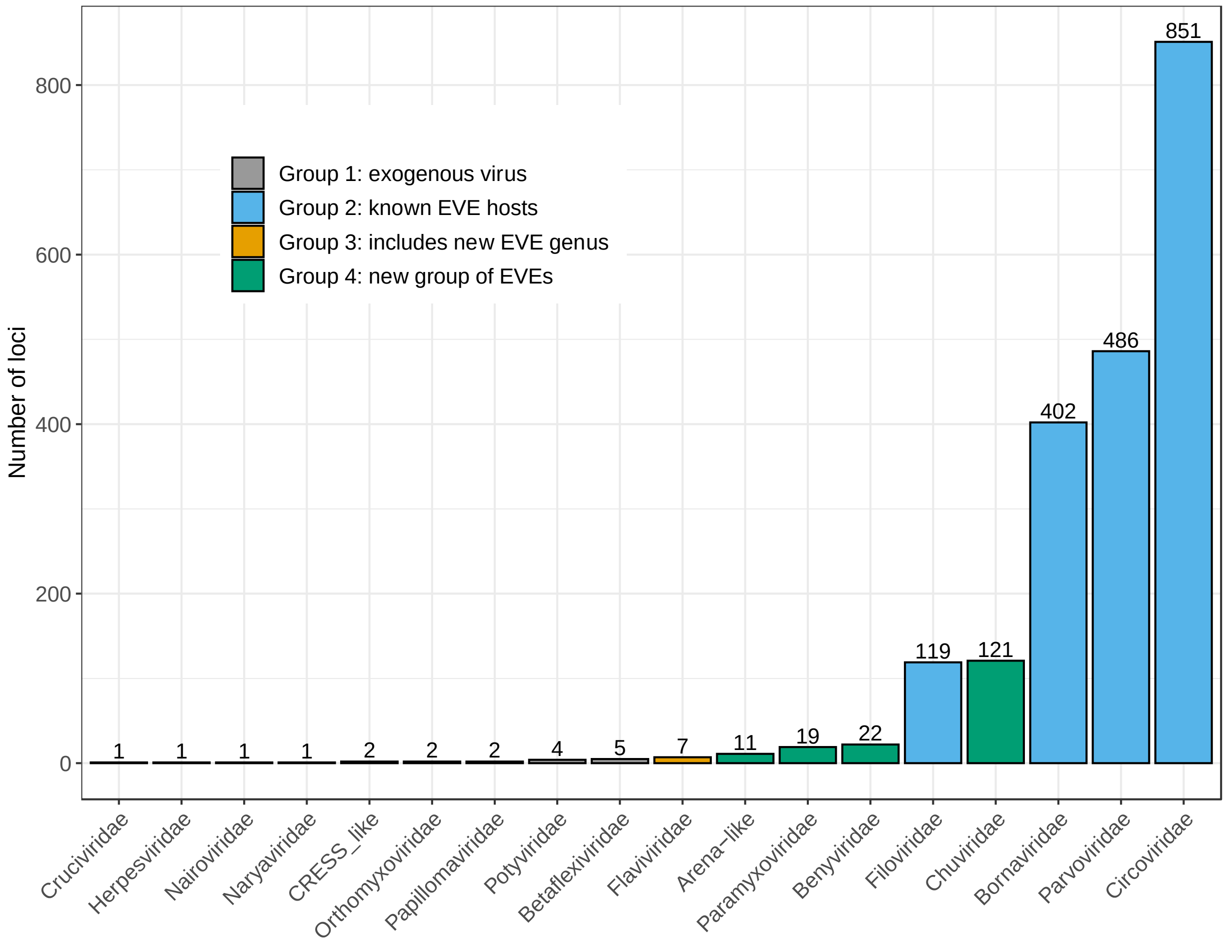


**Supplementary figure 1.** Distribution of non-redundant hits to viruses in representative vertebrate genomes (absolute numbers). In total 2,057 sequences were detected. Circovirus, parvovirus and bornavirus EVEs (1,739), account for 85.3% of all EVEs found (2,039). Sequences were divided into 4 groups depending on whether they are likely exogenous viruses (Group 1, grey), EVEs in known hosts (Group 2, blue), EVEs that include a new genus (Group 3, orange) or a new group of EVEs (Group 4, green). The “Arena-like” group refers to ectodomains with high similarity to reptarenavirus ectodomains, despite being embedded within retrovirus-like elements.

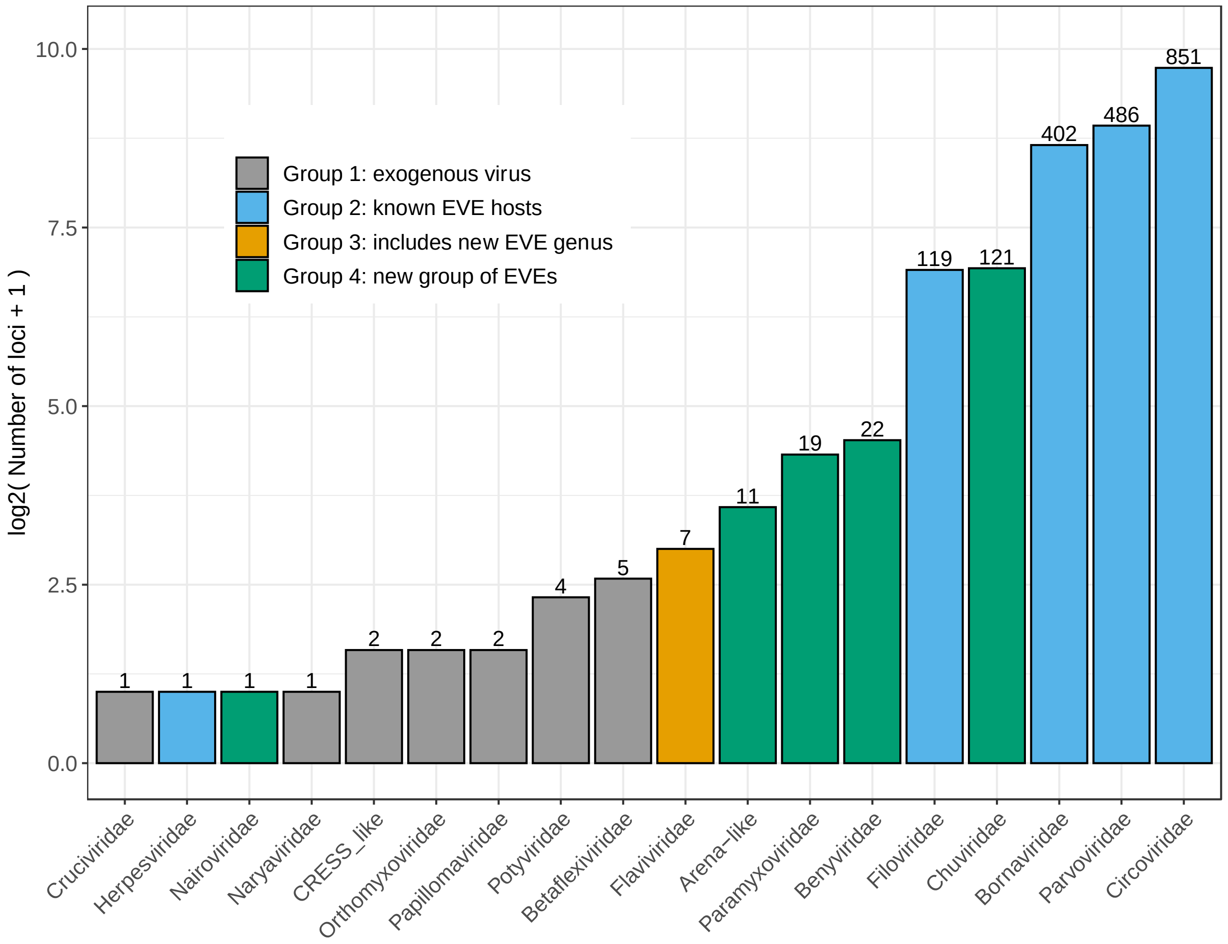


**Supplementary figure 2.** Distribution of non-redundant hits to viruses in representative vertebrate genomes (log2 scale). We used a log2-scale to facilitate comparisons of abundance. In total 2,057 sequences were detected. Circovirus, parvovirus and bornavirus EVEs (1,739), account for 85.3% of all EVEs found (2,039). Sequences were divided into 4 groups depending on whether they are likely exogenous viruses (Group 1, grey), EVEs in known hosts (Group 2, blue), EVEs that include a new genus (Group 3, orange) or a new group of EVEs (Group 4, green). The “Arena-like” group refers to ectodomains with high similarity to reptarenavirus ectodomains, despite being embedded within retrovirus-like elements.

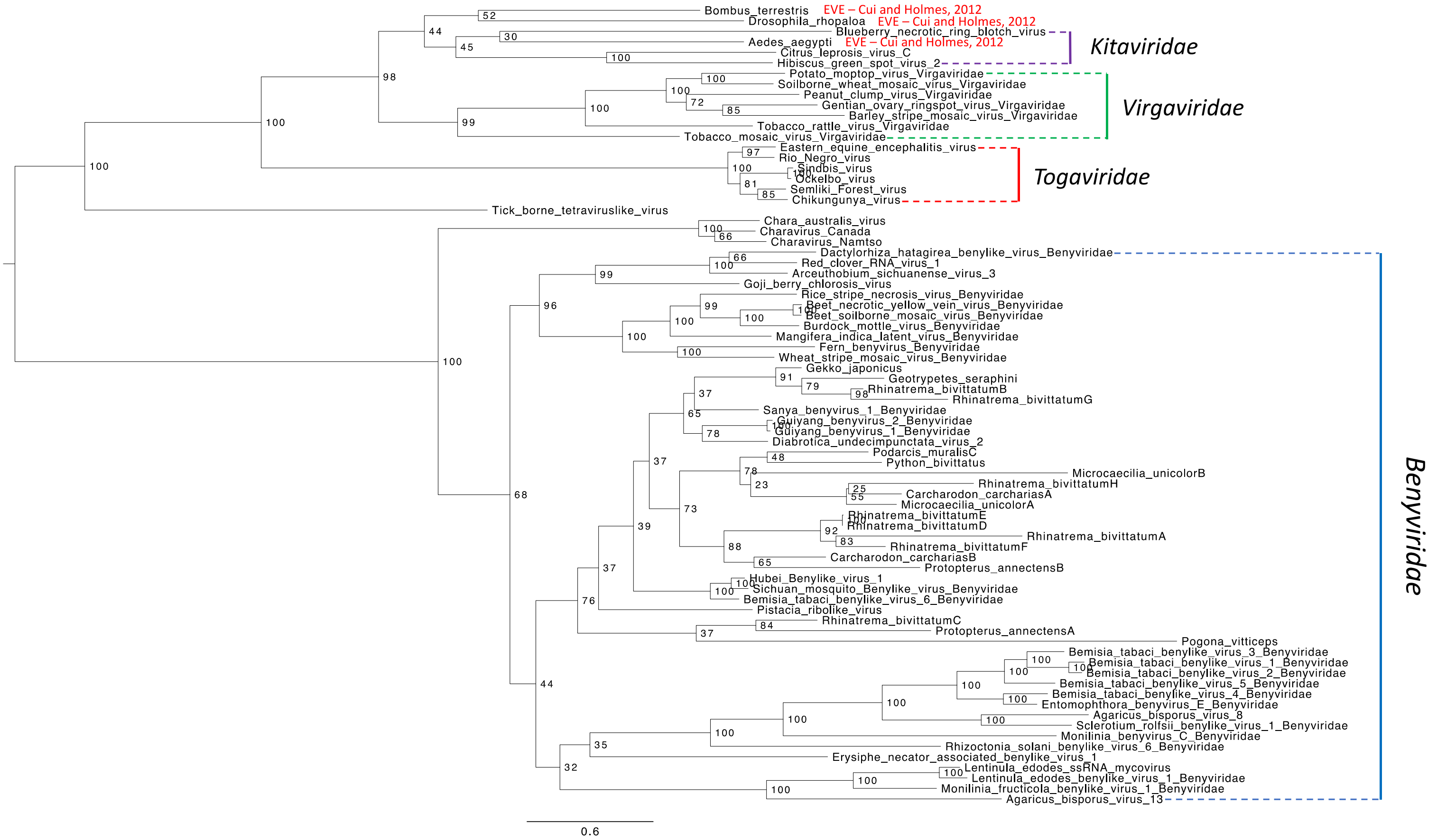


**Supplementary figure 3.** Midpoint-rooted maximum-likelihood tree of the RNA-dependent RNA polymerase of benyviruses, togaviruses, virgaviruses and kitaviruses, together with endogenous elements. Exogenous benyviruses and the vertebrate beny-like elements form a highly supported monophyletic group (100% bootstrap support) with charaviruses (which infect the charophyte alga *Chara*). Previously described plant virus-like EVEs in insect genomes (Cui and Holmes, 2012), fall closer to kitaviruses and virgaviruses. Tree inferred in RAxML-NG using the LG+FC+I+G4 model and 950 bootstrap replicates (until convergence).

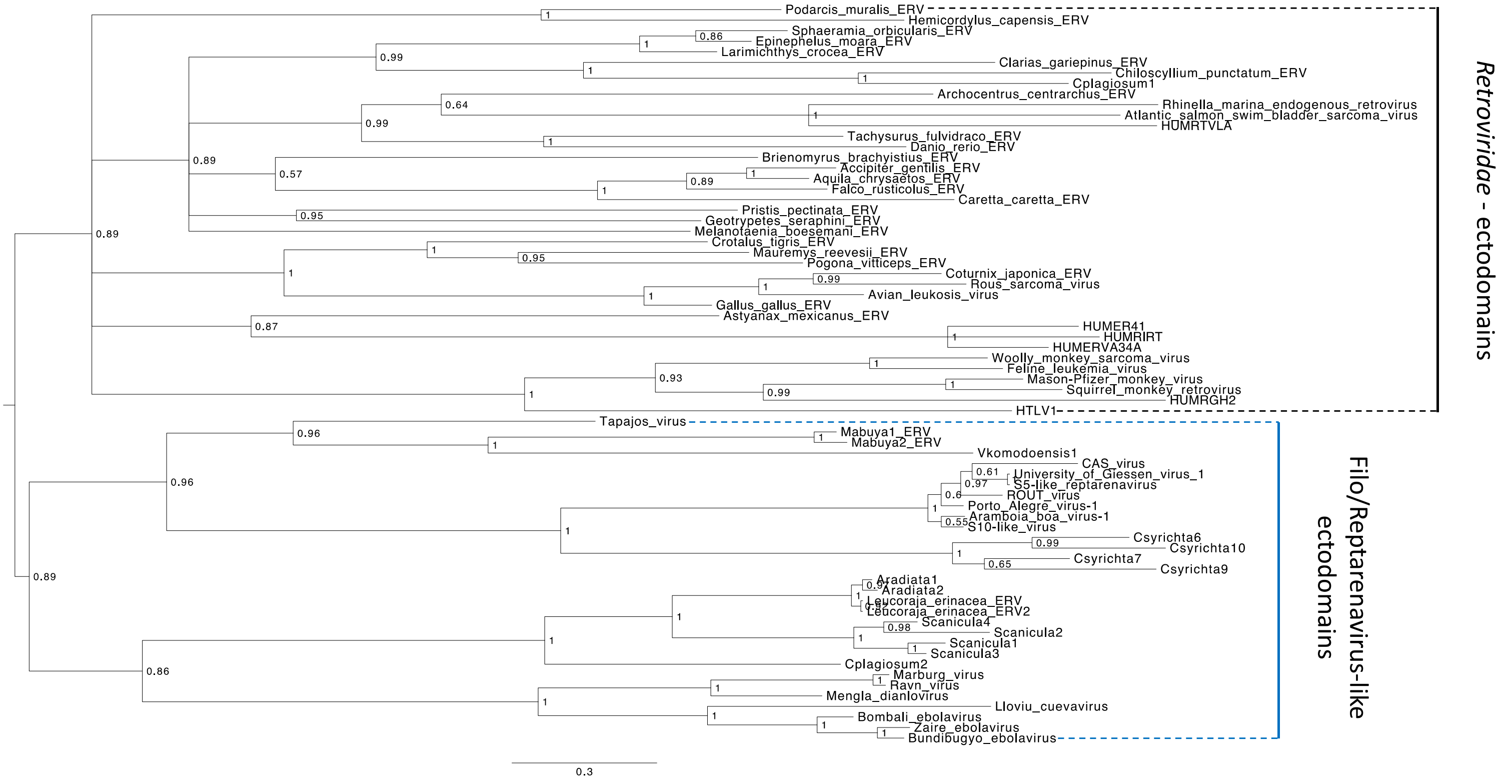


**Supplementary figure 4.** Midpoint-rooted Bayesian tree of ectodomains found in retroviruses, filoviruses, reptarenaviruses and endogenous elements in vertebrate genomes. The endogenous ectodomains found in the genomes of tarsiers (Csyrichta) and cartilaginous fish (Aradiata, Leucoraja, Scanicula, Cplagiosum) were placed in a clade (posterior probability = 0.89) together with the ectodomains of exogenous reptarenaviruses and filoviruses. Other retroviral sequences were placed outside this group. Tree inferred in MrBayes3 using the Vt+G4 model, 10 million generations and a 25% relative burn-in.

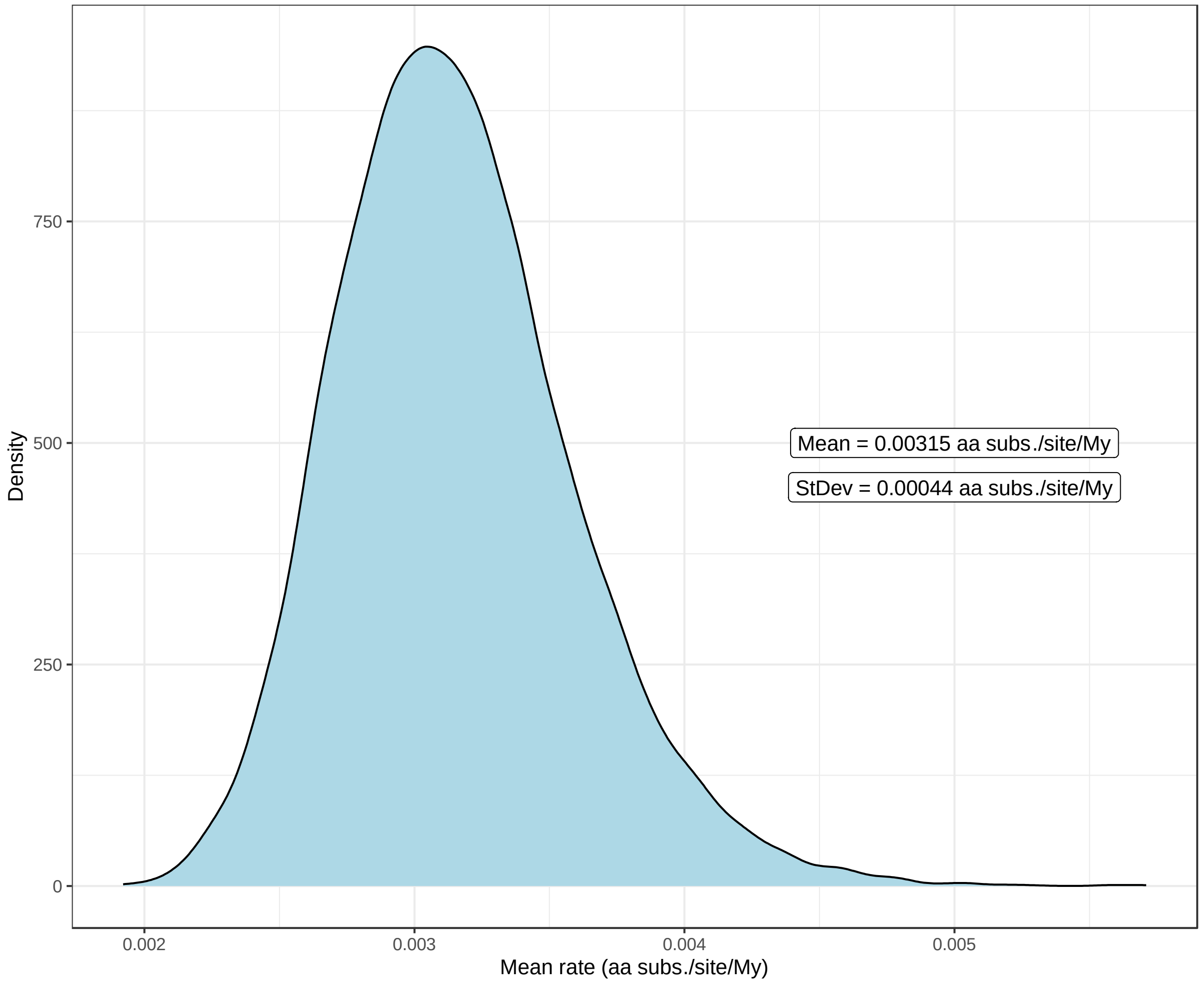


**Supplementary figure 5.** Distribution of mean evolutionary rates in the BEAST2 analysis. The mean evolutionary rate of the ectodomains was estimated at $3.2 \cdot 10^{-3}$ amino acid substitutions per site per million years (aa subs./site/My), with a standard deviation of $4.4 \cdot 10^{-4}$ (aa subs./site/My). Coefficient of variation = 0.14 (14%). Relative burn-in = 25%.
